# Efflux only impacts drug accumulation in actively growing cells

**DOI:** 10.1101/2021.05.11.443560

**Authors:** Emily E Whittle, Helen E McNeil, Eleftheria Trampari, Mark Webber, Tim W Overton, Jessica M A Blair

## Abstract

For antibiotics with intracellular targets, effective treatment of bacterial infections requires the drug to accumulate to a high concentration inside cells. Bacteria produce a complex cell envelope and possess drug-export efflux pumps to limit drug accumulation inside cells. Decreasing cell envelope permeability and increasing efflux pump activity can reduce intracellular accumulation of antibiotics, and are commonly seen in antibiotic resistant strains. Here, we show that the balance between influx and efflux differs depending on bacterial growth phase in Gramnegative bacteria. Accumulation of the model fluorescent drug, ethidium bromide (EtBr) was measured in *S*. Typhimurium SL1344 (wild-type) and efflux deficient (Δ*acrB*) strains during growth. In SL1344, EtBr accumulation remained low, regardless of growth phase and did not correlate with *acrAB* transcription. EtBr accumulation in Δ*acrB* was high in exponential phase but dropped sharply later in growth, with no significant difference to SL1344 in stationary phase. Low EtBr accumulation in stationary phase was not due to the upregulation of other efflux pumps, but instead, due to decreased permeability of the envelope in stationary phase. RNAseq identified changes in expression of several pathways that remodel the envelope in stationary phase, leading to lower permeability. This study shows that efflux is only important for maintaining low drug accumulation in actively growing cells, and that envelope permeability is the predominant factor dictating the rate of drug entry in stationary phase cells. This conclusion means that (i) antibiotics with intracellular targets may be less effective in complex non-growing or slow-growing bacterial infections where intracellular accumulation may be low, (ii) efflux inhibitors may be successful in potentiating the activity of existing antibiotics, but potentially only for bacterial infections where cells are actively growing and (iii) the remodelling of the cell envelope prior to stationary phase could provide novel drug targets.

## Introduction

Antibiotic treatment failure in clinical infections is increasingly common due to the rise in multi-drug resistant (MDR) Gram-negative bacteria. Gram-negative infections are particularly difficult to treat due to their impermeable outer membranes and efflux pumps which actively export antibiotic molecules out of the bacterial cell. Successful treatment relies on high concentrations of antibiotic accumulating within bacterial cells, which is a function of antibiotic influx and the rate of antibiotic efflux^1^.

Small hydrophilic antibiotics such as β-lactams enter a Gram-negative bacterial cell through membrane pores called porins. The major porins of *Enterobacteriaceae* are OmpF and OmpC^2^. Downregulation of porin genes contributes to antibiotic resistance by preventing antibiotics entering the cell^3^. In addition, mutations in the porin protein which change the channel diameter^4,5^ or the electric field inside the porin can block translocation of drugs across the membrane^5^.

Some drugs can enter Gram-negative cells through the lipid outer and inner membranes via ‘self-promoted uptake’. This mechanism has been described for EDTA, Polymyxin B, colistin and other cationic antimicrobial peptides (CAMPs), and aminoglycoside antibiotics^6–8^. The chelator, EDTA, acts as a permeabiliser by displacing and chelating the cations (Mg^2+^ or Ca^2+^) that are essential for the stability of LPS and the OM^6,9^. CAMPs interact with anionic groups on lipid A, breaching the outer membrane, and porate in the inner membrane, leading to bacterial death.

*Enterobacteriaceae* contain efflux pumps from 6 classes. MFS, SMR, MATE, RND and the recently described PACE pumps^10^ utilise the proton motive force for export of molecules such as antibiotics, and ABC (ATP binding cassette) pumps utilise ATP hydrolysis. Resistance-Nodulation-Division (RND) pumps are commonly upregulated in clinical isolates and can contribute to resistance to a number of antibiotic classes, as well as dyes, detergents and biocides^11^. The best described RND pump is AcrAB-TolC, found in *Salmonella enterica* serovar Typhimurium (*S*. Typhimurium) and *E. coli*. As efflux pumps underpin antibiotic resistance in essentially all bacteria of clinical and veterinary importance^12,13^ there is ongoing active research into the development of efflux inhibitors to potentiate the action of existing antibiotics.

Previous studies undertaken with cells in exponential growth phase have highlighted the importance of efflux pumps in minimising intracellular drug accumulation^14–17^. However, transcription of *acrAB* is growth phase dependent, with a peak in mid-exponential phase, which drops as cells enter into stationary phase^18^. The importance of AcrAB-TolC in bacterial cells in stationary phase which are slow-growing or non-growing is not known. However, it has been suggested that whereas survival of exponential-phase *E. coli* following treatment with the anionic detergent sodium dodecyl sulphate (SDS) is dependent on efflux, stationary phase cell survival is efflux-independent and rather is mediated by decreased permeability of the bacterial cell envelope, directed by the stationary phase sigma factor RpoS^19^. Little is known about the balance between influx and efflux in different growth phases and how this may relate to different growth states that may occur in an infection.

Previous studies have shown that the *E. coli* envelope changes in stationary phase when compared to logarithmic growth and it is possible that this could alter antibiotic influx in non-growing bacterial cells. Outer membrane changes include a decrease in the overall concentration of membrane proteins^20^ and an increase in lipoprotein crosslinked to peptidoglycan ^21,22^ to strengthen the outer barrier. In the inner membrane, the composition of fatty acids changes with a decrease in monounsaturated fatty acids^23^ and an increase in cyclopropane fatty acids, catalysed by Cfa^24^. Increased layers of peptidoglycan have also been described in stationary phase^25^.

Using a combination of fluorescent drug accumulation assays^17^, and measurement of efflux gene transcription in wild-type and efflux mutant strains we here assess the importance of the balance between influx and efflux in different growth phases in Gram-negative bacteria, using the model organism *Salmonella enterica* serovar Typhimurium. We also use RNASeq to measure the global transcriptome as bacteria enter stationary phase and correlate transcriptomic changes with biochemical and physiological changes in the cell envelope that lead to alterations in permeability.

## Results

### Accumulation level of drugs by *S*. Typhimurium is independent of growth phase-dependent *acrAB* transcription

Using a recently developed flow cytometry method^17^, both the intracellular accumulation of the fluorescent dye ethidium bromide (EtBr), and the transcription of *acrAB* (via a promoter-GFP fusion) were measured in parallel in single cells of *Salmonella* grown in *drug free* media. Samples were taken hourly during batch culture before EtBr was added to the sample immediately prior to flow cytometry analysis to measure accumulation (EtBr was *not* present in growing the culture).

Transcription of *acrAB* in SL1344 was growth phase dependent and peaked in early-mid log phase before decreasing towards stationary phase (**Figure 1**), as previously described^18^. Previous studies have shown that increased expression of *acrAB* in clinical isolates leads to decreased susceptibility to antibiotics^12^. Given the known role of efflux pumps in drug export, one might predict that EtBr accumulation would be lowest when efflux expression was highest. Our data however show that this is not the case. In SL1344 cells, accumulation of EtBr was low and remained unchanged across growth despite changes in *acrAB* transcription (Fig 1). Therefore, changes in efflux pump transcription in different growth phases does not alter levels of drug accumulation within the cell.

**Figure 1.**
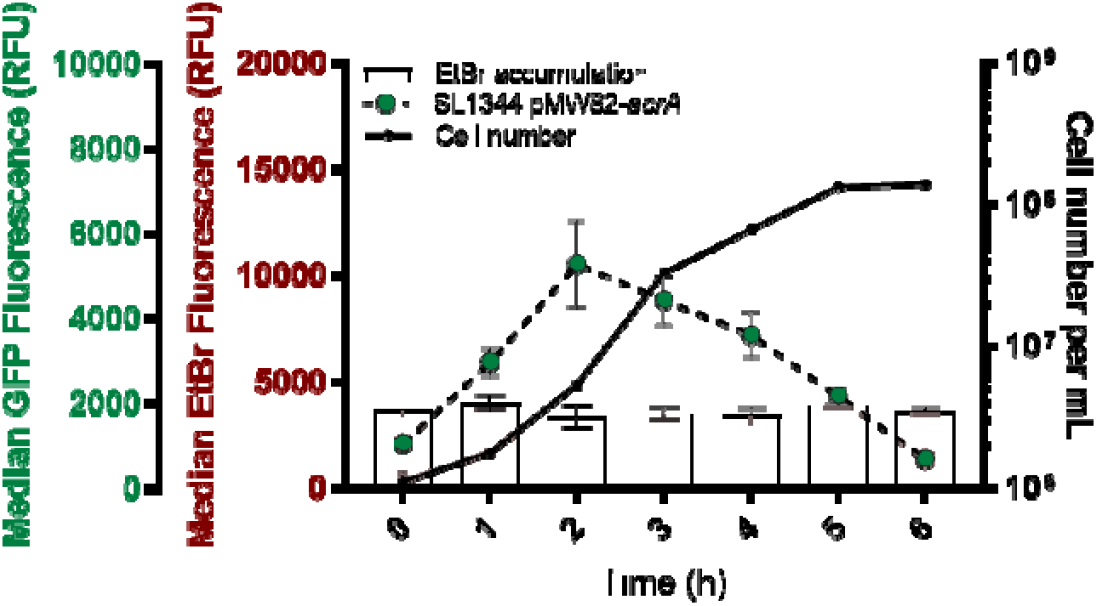
Ethidium bromide accumulation and *acrAB* expression in single cells of *S*. Typhimurium SL1344 across the growth phase. Cell number per mL was measured in each sample (black lines, numbers indicated on the right y-axis). Pink bars indicate median ethidium bromide fluorescence per cell (relating to left red y-axis) and dashed lines with green circles show *acrAB* expression (median GFP fluorescence per cell from reporter, relating to left green y-axis. All data points are median values from measurements of 10,000 single cells of SL1344. Error bars indicate standard error of the mean (+/- SEM).

### Growth phase-dependent transcription of *acrAB* does not correlate with drug accumulation, efflux capacity or AcrAB protein level

Having shown that *acrAB* transcription does not correlate with ethidium bromide accumulation, the efflux function in a population of cells was measured to determine whether efflux activity varied with growth phase (and *acrAB* expression), even if drug accumulation did not.

To measure functional efflux capacity of cells we used the previously described direct efflux activity assay^14^ which was further optimised to analyse efflux capacity at 3 different time points across growth in SL1344. This assay determines the efflux capacity of the cell based on the activity of all efflux pumps (not just AcrAB-TolC) that are able to transport EtBr. Cultures grown for 1, 3 and 5 hours had the same capacity to efflux the substrate as there was no significant difference in efflux rate between samples taken at each time point (**Figure 2A**) (based on time taken for ethidium bromide fluorescence to drop 10%, 25% and 50% from its maximum fluorescence value) regardless of the different levels of *acrAB* transcription at these time points already established.

**Figure 2.**
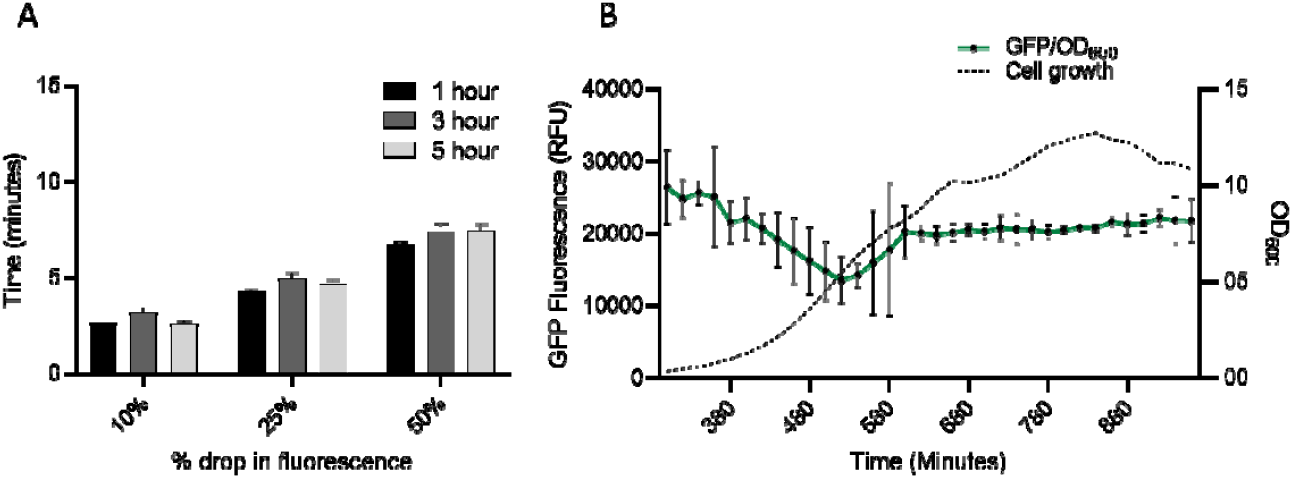
(A) Time taken for ethidium bromide to be removed from SL1344 cells at 1,3 and 5 hours. Bars represent the time taken for ethidium bromide fluorescence to drop 10%, 25% and 50% from its original value. Data is based on 3 biological replicates with error bars showing standard error of the means (SEM). 1 hour (black), 3 hours (dark grey) and 5 hours (light grey) are shown. There was no significant difference in the time taken to export EtBr at each time point. **(B) GFP/OD_600_ from SL1344 AcrB-GFP over 16 hours of growth in MOPs minimal media.** This graph shows GFP/OD_600_ from AcrB-GFP at the end of lag phase (300 minutes) until the last time point at 16 hours. The dashed black line shows the OD_600_ whereas the green line error bars +/- SEM shows GFP fluorescence. SL1344 autofluorescence was subtracted from this data.

Taken together, the low accumulation and similar rate of efflux of EtBr across time in SL1344 suggests that although *acrAB* transcription peaks in mid-exponential phase, activity of the assembled AcrAB-TolC complex remains constant. The AcrB protein is known to be very stable once made with a predicted half-life of 6 days^26^. To measure AcrB protein level at different points during growth a strain was constructed in which the AcrB protein was tagged with GFP at the C-terminus as previously described ^27^. The generation time and efflux level in this strain were unaffected confirming that tagging GFP to the C-terminus of AcrB did not affect its function. Measurement of GFP fluorescence during 16 hours of growth showed that AcrB level remains constant (**Figure 2B**). These data suggest that efflux capacity is constant regardless of growth phase due to the constant level of AcrAB protein within a population and may explain why EtBr accumulation remained low in stationary phase despite decreased efflux gene transcription.

### Drug accumulation is only dependent on efflux in actively growing cells

To further dissect the importance of efflux during different growth stages we measured EtBr accumulation (as in Fig. 1) in the presence or absence of AcrAB-TolC function (using SL1344 Δ*acrB*). The previous results suggest AcrAB-TolC activity is constant, therefore by removing the efflux pump, it was assumed that EtBr accumulation would be high across growth. When measuring EtBr accumulation in SL1344 Δ*acrB*, after 1 hour of growth, EtBr accumulation was 6-fold higher than in SL1344. This is similar to the growth time point used in most other published studies that have shown an increase in accumulation upon deletion of *acrB*^11,14,15,17^. However, EtBr accumulation then decreased dramatically and was not significantly different from WT from 3-6 hours of growth (**Figure 3A**). This suggests that low accumulation at 1 hour in SL1344 greatly depends on efflux to export ethidium bromide from actively growing cells. As there is no significant difference between Δ*acrB* and WT cells from 3-6 hours, it suggests that AcrAB-TolC is not important in maintaining low accumulation in slower growing or stationary phase cells. This is also supported by the *acrAB* expression data which shows highest expression in the early stages of logarithmic growth.

**Figure 3.**
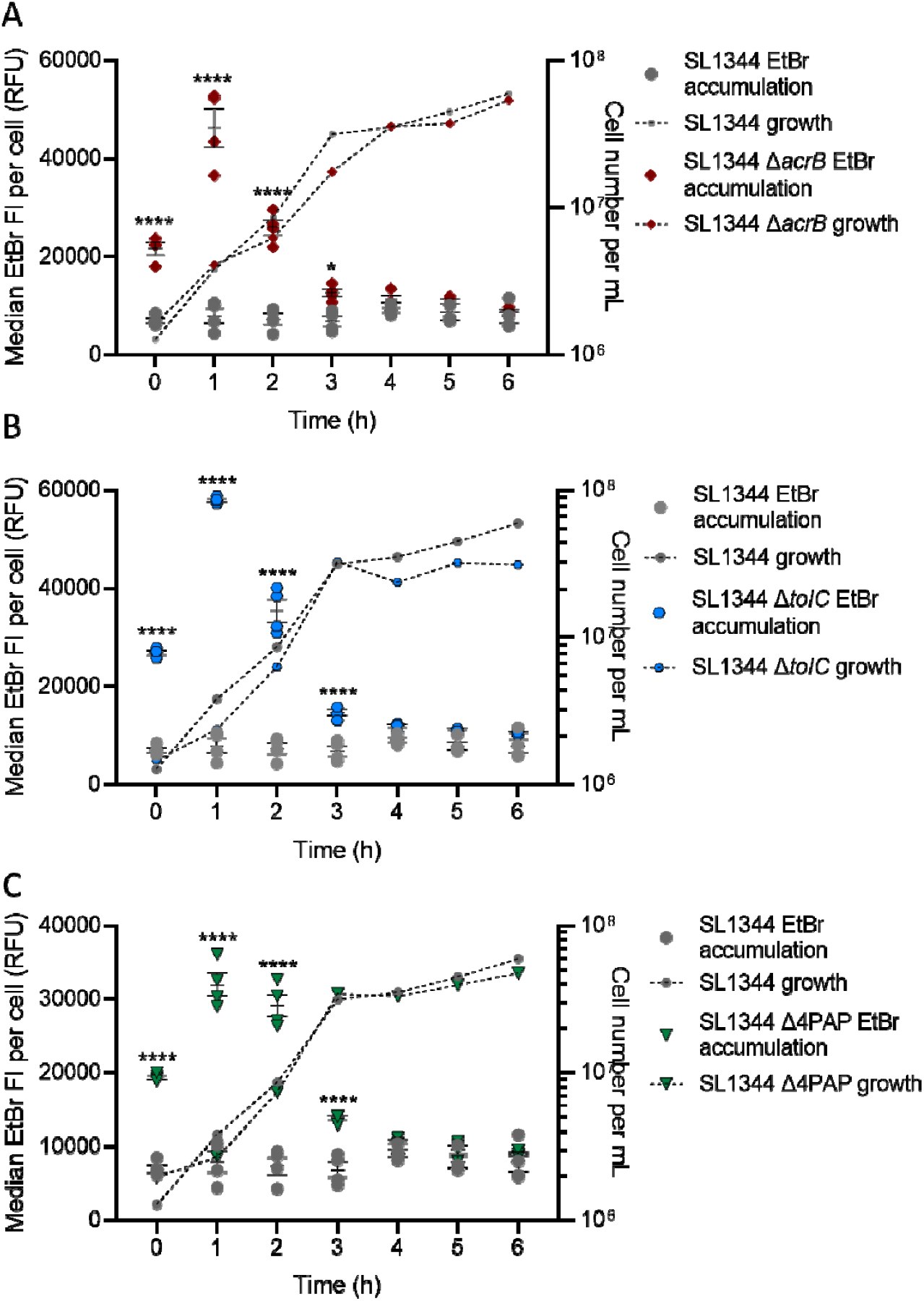
EtBr accumulation in SL1344 and SL1344 Δ*acrB* (A), Δ*tolC* (B) and Δ4PAP (C). For each strain the median EtBr fluorescence per cell in 10,000 single cells was measured every hour between 0 and 6 hours of growth for SL1344 (Grey circles) and (A) SL1344 *ΔacrB* (red diamonds) (B) Δ*tolC* (blue hexagons) and (C) Δ4PAP (Δ*acrA* Δ*acrE* Δ*mdsA* Δ*mdtA*) (green triangles). Data from 4 biological replicates for each strain are shown, horizontal bars show the mean and error bars the SEM. Median EtBr fluorescence per cell is plotted on the left y-axis. Calculated cell number per mL values were plotted on the right y-axis with corresponding symbols equating to strain and a dashed line to show growth of the cultures. Cell numbers were based on the mean of the same biological replicates and the same gated population that EtBr fluorescence was measured from. Two-way ANOVA and Sidak’s multiple comparison test were carried out for statistical analysis. At 0, 1 and 2 hours, EtBr accumulation was significantly increased in Δ*acrB* with p values of <0.0001 (****). At 0, 1, 2 and 3 hours, EtBr accumulation was significantly increased in Δ*tolC* and Δ4PAP with p values of <0.0001.

To confirm that low EtBr accumulation in stationary phase was not due to the activity of other RND efflux pumps present in SL1344, EtBr accumulation was also measured in two other mutants of SL1344. In the first, *tolC* was deleted which compromises most efflux systems in *Salmonella* which use TolC as a common outer membrane channel. The second strain used lacked all four periplasmic adaptor proteins (Δ4PAP; Δ*acrA* Δ*acrE* Δ*mdsA* Δ*mdtA*) and is incapable of assembling any functional RND efflux systems. In both strains, the EtBr accumulation pattern observed recapitulated that seen in SL1344 Δ*acrB*, with a peak in accumulation at 1 hour, but no significant difference to SL1344 in stationary phase cells (**Figure 3B&C**). This result showed that low accumulation in stationary phase was not due to any RND pump in SL1344, (nor the ABC pump MacAB-TolC). In addition, we also showed that, apart from *acrAB* whose transcription was highest in mid-log phase and lowest in stationary phase, no other RND pump was actively transcribed in the conditions used to measure accumulation capacity across growth (**Figure S1**). For pumps from other families, only *macA* (ABC), *mdfA* (MFS) and *mdtK* (MATE) were transcribed and only at low levels (**Figure S1**).

Further investigation into the role of efflux pumps in stationary phase EtBr accumulation was carried out by measurement in the presence of the proton motive force inhibitor CCCP. Inhibiting the proton motive force, inhibits the activity of the RND, MFS and MATE pumps of SL1344^28,29^. In SL1344 in the presence of CCCP, EtBr accumulation peaked at 1 hour (**Figure S2**). Accumulation levels started to drop into stationary phase, strikingly similar to SL1344 Δ*acrB*, again suggesting that low accumulation in stationary phase is not dependent on RND, MFS or MATE-mediated efflux. This independent confirmation using different mutants and inhibitors demonstrates that the observed low EtBr accumulation in stationary phase is efflux-independent.

To investigate whether this was just a *Salmonella* phenomenon, EtBr accumulation was measured in wild-type and a mutant lacking major RND efflux pump of other Gram-negative bacterial species including *Escherichia coli* (MG1655 and MG1655 Δ*acrB*), *Pseudomonas aeruginosa* (PA01 and PA01 Δ*mexA*) and *Klebsiella pneumoniae* (ecl8 and ecl8 *acrB*::Gm). In *E. coli* and *K. pneumoniae*, EtBr accumulation was low throughout growth for the wild-type but peaked at 1 hour for each *acrB* mutant (**Figure S3**) and in *P. aeruginosa*, the *mexA* mutant peaked at 2 hours (**Figure S4**) and then dropped to WT levels in stationary phase. Therefore, very similar observations are seen in a wide range of Gram-negative organisms.

The EtBr accumulation pattern in *Salmonella* was also shown in MOPS minimal medium suggesting that the pattern was not influenced by media type and specifically was not a result of the limitations of LB^30^ (**Figure S5**). Even though is a well-established and studied model efflux substrate, to counter the possibility that EtBr would give abnormal results which are not representative of other efflux substrates, the same accumulation pattern in *Salmonella* WT and Δ*tolC* was also shown using the lipophilic dye Nile Red (**Figure S6**). Unlike EtBr, Nile Red does not fluoresce on intercalation with DNA, but rather when bound to phospholipids or triglycerides^31^ showing this is not an artefact of the dye initially used. Together this data shows that the accumulation pattern described in the absence of efflux is consistent regardless of Gram-negative species, media type or efflux substrate used.

### Drug accumulation in stationary phase is controlled by reduced membrane permeability

Together this data shows that cells from later growth phases minimise intracellular accumulation of EtBr (and other substrates) in an efflux independent manner. We hypothesised this could be due to a shift in the balance between influx and efflux over growth, with influx rate, controlled by reduced permeability of the outer membrane, being more important in slower growing or stationary phase cells.

Several dyes that are often used to probe the permeability of the outer membrane, such as NPN (1-*N*-Phenylnaphthylamine), are efflux substrates and therefore assessing membrane permeability in strains lacking efflux pumps is problematic. Most hydrophilic antibiotics enter Gram-negative bacterial cells through outer membrane porins such as OmpC and OmpF. To investigate whether porins altered the accumulation of EtBr, accumulation assays were performed using SL1344 mutants; Δ*ompC*/Δ*ompF*/Δ*acrB*, Δ*ompC*/Δ*acrB*, Δ*ompF*/Δ*acrB*, Δ*ompC*, and Δ*ompF* and showed that none had a significantly different EtBr accumulation pattern to those previously seen, confirming that EtBr doesn’t enter *S*. Typhimurium through OmpF or OmpC (**Figure S7**). A similar observation was made by Murata *et al*. in *E. coli* K-12 ^32^, who concluded that the OM bilayer is the predominant mode of EtBr entry.

Since SYTO 84 is used in our flow cytometry assay as a probe to stain cells, the accumulation of this dye was first investigated to assess permeability and to determine if it is an efflux substrate. SYTO 84 is marketed as a cell-permeant DNA dye and so is expected to readily enter bacteria. There was no significant difference between the accumulation of SYTO 84 in SL1344 and SL1344 Δ*acrB* and in both strains accumulation peaked after 1 hour of growth (**Figure 4**). This shows that SYTO 84 is not an efflux substrate and demonstrates the importance of efflux in maintaining low accumulation of drugs and dyes that are substrates in actively growing cells. However, SYTO 84 fluorescence decreased significantly in both strains on entrance to stationary phase. This suggests that a compound that is not exported via efflux, is also less able to enter bacteria during stationary phase and we hypothesise this is due to a strengthening of the permeability barrier. It is important to note that, although the SYTO 84 fluorescence does reduce around 2.5-fold in stationary phase, the lowest value is still over 45,000 RFU, so the reduction does not compromise its use to differentiate cells from acellular particles in the EtBr accumulation assays using flow cytometry.

**Figure 4.**
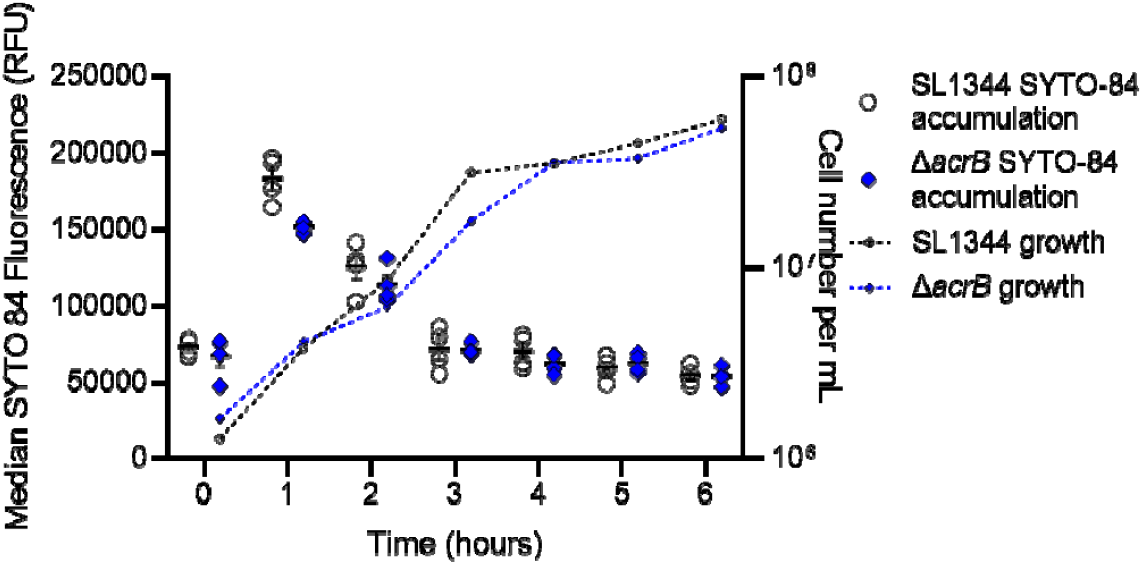
SYTO 84 accumulation in SL1344 and SL1344 Δ*acrB*. Median SYTO 84 fluorescence per cell in 10,000 cells was measured every hour between 0 and 6 hours. White circles (SL1344) and blue diamonds (Δ*acrB*) represent the X-median value of SYTO 84 fluorescence in 10,000 cells within a biological replicate. 4 biological replicates for each strain are shown, with +/- SEM error bars. Median SYTO 84 fluorescence is plotted on the left Y-axis. Calculated cell number per mL values were plotted on the right Y-axis with corresponding symbols equating to strain and a dashed line to show growth of the culture. Cell numbers were based on the mean of the same biological replicates and the same gated population that EtBr fluorescence was measured from.

Ethidium bromide is a cationic dye that diffuses into cells through the OM^32^. LPS molecules on the outer face of the outer membrane are ionically cross-linked to each other by divalent cations (Mg^2+^ or Ca^2+^) binding to phosphate groups in lipid A, generating a permeability barrier. EDTA is considered a ‘permeabiliser’ which can chelate and thus displace divalent cations, destabilising and releasing LPS^33^, thereby increasing the permeability of the cell to itself and other compounds^6^. Increasing concentrations of EDTA were used to permeabilise the outer membrane and assess the effect on ethidium bromide accumulation (**Figure 5A**). Following 1 or 3 hours of growth, there was no significant difference in EtBr accumulation up to 100 μM EDTA. At 200 μM and 500 μM EDTA, EtBr accumulation was significantly higher, suggesting that EDTA was able to make the outer membrane more permeable to EtBr. At 5 hours, neither 200 μM nor 500 μM ETDA had any effect on the accumulation of EtBr. This suggests that the *Salmonella* outer membrane is remodelled during entry into stationary phase and becomes less reliant on cation-mediated crosslinking to maintain its permeability barrier to EtBr. Indeed, both *Salmonella* and *E. coli* become more resistant to CAMPs, whose mode of action relies upon interaction with negative charges on the LPS, in stationary phase^34,35^

**Figure 5.**
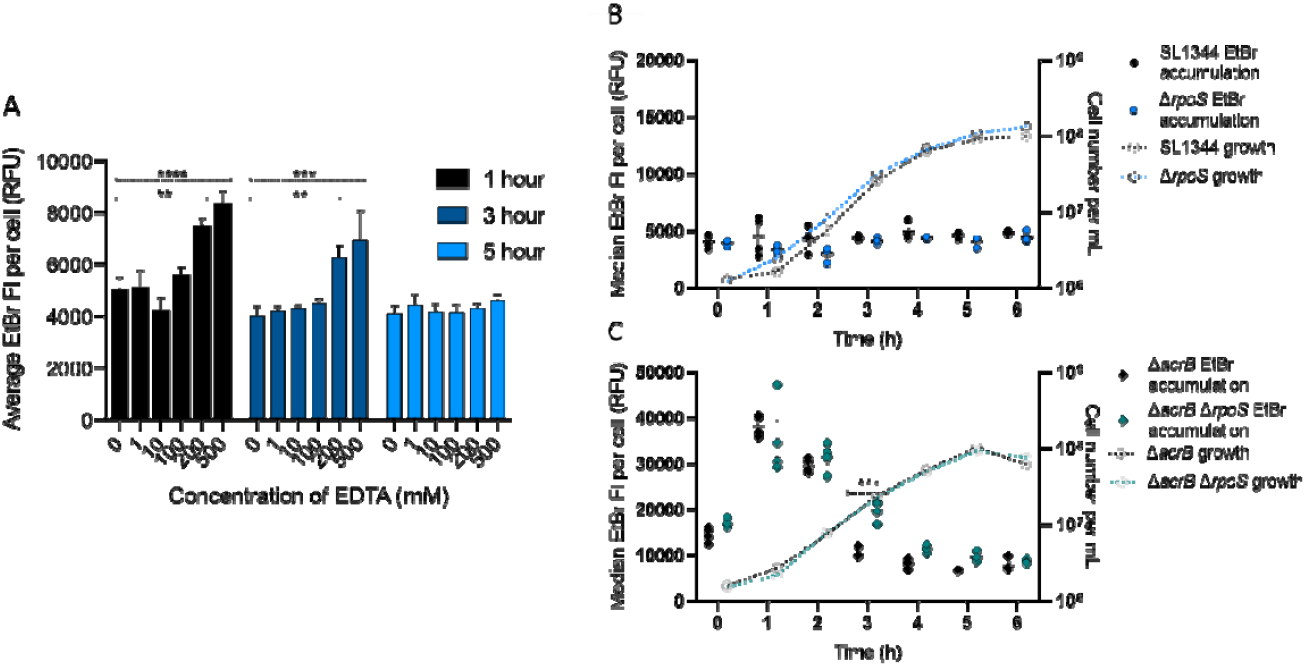
**(A) EtBr accumulation in SL1344 treated with EDTA.** Bars represent median EtBr fluorescence in 10,000 single cells of SL1344. EtBr accumulation was measured in the presence of increasing concentrations of EDTA (0, 1, 10, 100, 200 and 500 mM) from a culture grown for 1 hour (black), 3 hours (dark blue) and 5 hours (light blue). Error bars show SEM from 3 biological replicates. Dashed lines above the bars with asterisks represent significance value when based on a T-test compared to no EDTA added. At 1 hour, treatment with 200 and 500 mM significantly increased EtBr accumulation in SL1344 with p values of 0.0013 (**) and <0.0001 (****) respectively. At 3 hours, treatment with 200 and 500 mM significantly increased EtBr accumulation in SL1344 with p values of 0.0033 (**) and 0.0001 (***) respectively. **(B+C) EtBr accumulation in SL1344 Δ*rpoS* and Δ*acrB* Δ*rpoS*.** 4 biological replicates for each strain are shown, with a short mean bar and SEM error bars. EtBr accumulation is plotted on the left Y-axis. Calculated cell number values were plotted on right Y-axis. Cell numbers were based on the mean of the same biological replicates and the same gated population that EtBr fluorescence was measured from. **(B)** shows SL1344 WT (individual black dots) vs Δ*rpoS* (blue dots). Median EtBr fluorescence per cell in 10,000 SYTO-84^+^ flow cytometry events was measured every hour between 0 and 6 hours. Individual symbols represent the median value of EtBr fluorescence within a biological replicate. **(C)** shows SL1344 Δ*acrB* (black diamonds) vs Δ*acrB ΔrpoS* (green diamonds). Median EtBr fluorescence per cell in 10,000 SYTO-84^+^ flow cytometry events was measured every hour between 0 and 6 hours. Individual symbols represent the median value of EtBr fluorescence within a biological replicate. Significant differences to parent strain were measured by a two-way ANOVA and Sidak’s multiple comparison test. At 3 hours, EtBr accumulation in Δ*acrB ΔrpoS* is significantly different to Δ*acrB* with a p value of 0.0002 (***).

A previous study found that increased SDS resistance in carbon-limited stationary phase *E. coli* is due to decreased envelope permeability mediated by RpoS-dependent and −independent mechanisms^19^. The role of RpoS in decreased EtBr permeability in *S*. Typhimurium was therefore investigated by construction of Δ*rpoS* mutants of SL1344 and its Δ*acrB* variant.

Deletion of *rpoS* in SL1344 caused no significant difference in EtBr accumulation (**Figure 5B**), although these bacteria were efflux-active so EtBr could be pumped out. Comparison of the Δ*acrB* and Δ*rpoS ΔacrB* mutants (**Figure 5C**) revealed a significant difference in EtBr accumulation only around 3 h growth; the Δ*rpoS* mutant showed a delayed decrease in EtBr accumulation, although in stationary phase the two strains were similar. We conclude that in *S*. Typhimurium, although RpoS might play a role in envelope remodelling, it is not essential for generation of a low-permeability envelope in stationary phase, so there are likely to be RpoS-dependent and −independent pathways to achieve this phenotype. Although SDS and EDTA disrupt the cell envelope in different ways (detergent disruption of lipid membranes versus chelation of divalent cations), it is clear that RpoS-dependent and – independent mechanisms play a role in envelope remodelling in both *E. coli*^19^ and *S*. Typhimurium.

### RNAseq analysis identified several pathways likely to be involved in reduced envelope permeability in S. Typhimurium

Given the data above did not identify a definitive mechanism by which the stationary phase cell envelope displays lower permeability to EtBr, we used RNAseq analysis to identify genes and pathways that may be involved in changes to Gram-negative cells as they enter stationary phase. Growing cultures of SL1344 were sampled after 1 hour, 3 hours and 5 hours of growth and RNA was extracted and analysed by GENEWIZ Inc. Comparing SL1344 at 1 hour versus 3 or 5 hours of growth, 1228 (26%) and 2260 (47%) genes were differentially expressed respectively. The data is deposited with Array Express (Accession: E-MTAB-9679). Differentially-expressed genes were then identified that encode proteins involved in envelope remodeling in stationary phase, many of which have been shown to increase barrier function (Supplementary Table S1, summarized in Figure 6).

**Figure 6.**
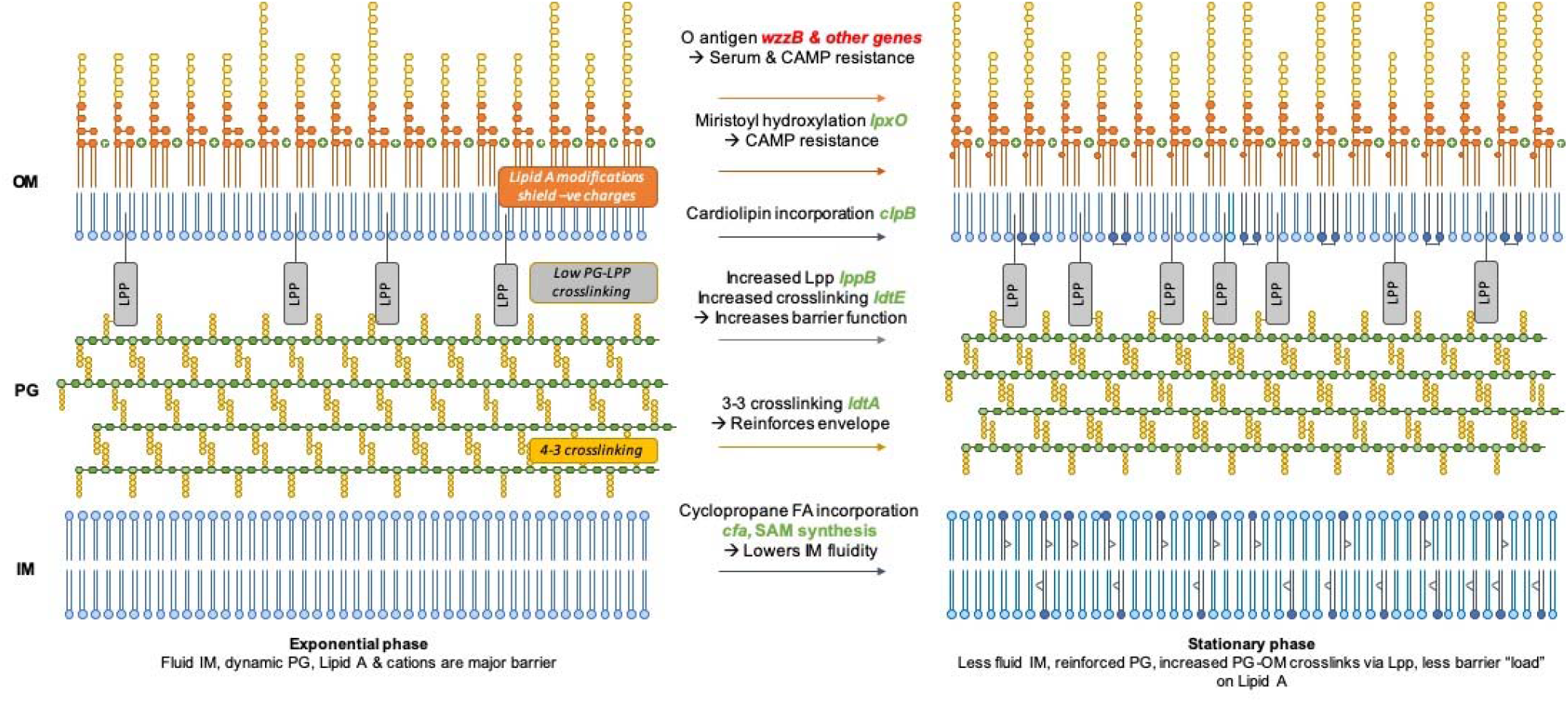
Model showing that the differentially-expressed genes identified in the RNAseq encode proteins involved in envelope remodeling in stationary phase to increase barrier function.

Previous studies have suggested that multiple layers of the cell envelope are remodeled upon entry into stationary phase^34^ and our RNASeq data support this; full description of this dataset is in the supplementary material. Inner membrane fluidity decreases with cyclopropane fatty acid incorporation^36–38^, mediated by upregulation of *cfa*. Stationary phase peptidoglycan contains 3-3 (LD) rather than 4-3 (DD) crosslinks^22,39,40^; relevant transpeptidases are up- and downregulated. The quantity of Lpp in the OM increases (*lppB* is upregulated) and becomes more highly crosslinked to the PG (*ldtE* is upregulated^22^), which has been shown to increase barrier function^41^. OM inner leaflet cardiolipin content is known to increase^42^ potentially mediated by upregulation of *clsB*. LPS modification pathways important in exponential phase (e.g. the *pmr* genes which confer CAMP resistance primarily through negative charge neutralization)^43–45^ are downregulated whereas *lpxO* (involved in myristoyl chain hydroxylation and implicated in CAMP resistance in *K. pneumonia*^46^) is upregulated. Genes involved in O-antigen synthesis and chain length regulation are downregulated; average O-antigen chain length increases in stationary phase and O-antigen structure has been shown to influence serum resistance^47^ and CAMP susceptibility^48^. Finally, genes involved in enterobacterial common antigen (ECA) synthesis are downregulated; ECA is implicated in envelope integrity and bile resistance.

Taken together, this leads to a model (Figure 6) suggesting why the exponential phase cell envelope is more susceptible to attack from various factors (EtBr, CAMPs, EDTA, antibiotics). Resistance to self-mediated uptake in exponential phase is provided primarily by the barrier function of LPS, comprising lipid hydrophobicity and crosslinking between phosphate groups and divalent cations. The LPS takes the burden because the inner layers of the envelope (PG and IM) are by necessity more fluid; PG is being extensively and continually remodeled to permit growth and division, and the IM is similarly fluid. The reliance on the LPS as the primary barrier poses problems when antimicrobials such as EDTA and CAMPs target the phosphate-cation bridges. The cell responds by shielding negative charges and modifying the LPS lipid content to decrease fluidity, regulated by PmrAB.

In stationary phase, each layer of the envelope plays a greater role in barrier function because remodeling and fluidity is less of a requirement. The lipid components of the IM and OM become less fluid and the PG contains more LD-crosslinks and becomes more crosslinked to the OM, further strengthening the OM permeability barrier and decreasing (but not eliminating) the requirement for cation crosslinking of LPS. The saccharide components of the LPS provide the outermost layer of protection. This “laminated” approach shares the burden of protection and generates a strong barrier against multiple chemicals which seek to enter and damage the cell, reflected by the increased resistance of stationary phase cells to multiple stressors.

## Discussion

This study shows that the mechanisms that control drug accumulation are growth phase dependent. In actively growing cells, efflux is fundamental to maintaining low drug accumulation and subsequently survival of the bacterial population. Bacterial infections are complex, and bacterial populations will often not be in a single growth phase, therefore more careful consideration may be required for the most effective antibiotic treatment. We have shown that stationary phase slow or non-growing cells are impermeable, and that this is not due to changes in porin production but as a result of membrane remodeling and increased peptidoglycan crosslinking which reinforces the envelope barrier function. Treatment of chronic infections and biofilms where bacterial cells are slow- or non-growing may need to be considered more carefully. Successful treatment of these infections is already extremely difficult, and careful consideration is already made for treatment of intrinsically impermeable pathogens such as *Pseudomonas aeruginosa* and *Acinetobacter baumanii*. More extensive research into the effects of an impermeable membrane on treatment during infection must now be carried out.

Efflux pumps are only important in maintaining low drug accumulation in actively growing cells which have a more permeable envelope. If an infection is actively growing, it seems likely that efflux inhibitors would be effective at increasing the accumulation of antibiotics within cells to potentiate their activity. However, if cells are in a slow-growing or non-growing state, where membrane permeability is fundamental to maintaining low drug accumulation, efflux inhibitors may not be an effective treatment option. It is also possible that administering an efflux inhibitor where it has no effect on treating an infection, may also lead to the development of new mechanisms of AMR.

## Materials and Methods

### Strains and growth conditions

Unless otherwise stated, all experiments use *Salmonella enterica* serovar Typhimurium (hereafter named *S*. Typhimurium^49^) SL1344. The Δ*acrB* and Δ4PAP strains (Δ*acrA* Δ*acrE* Δ*mdsA* Δ*mdtA*) have been previously published^50,51^. SL1344 Δ*ompF* and Δ*ompC* strains were constructed for this study using the Datsenko and Wanner method of gene deletion^52^. Transcriptional reporter constructs were made by fusing the promoter of each efflux pump gene to *gfp* in the pMW82 plasmid^53^. These plasmids were transformed into SL1344 and SL1344 Δ*acrB. E. coli* MG1655 Δ*acrB*^15^, *P. aeruginosa* PA01 Δ*mexA*^54^ and *K. pneumoniae* ecl8 *acrB::Gm*^17^ were also used as part of this study and are previously published.

Unless otherwise stated, LB (Sigma) was used as growth medium for all assays. One assay used MOPs minimal media (Teknova) which was supplemented with 400 mg/L histidine.

### Chromosomal insertion of *gfp* downstream of *acrB* to produce SL1344 AcrB-GFP

To measure the protein level of AcrB in *S*. Typhimurium, a gene encoding a monomeric super-folder GFP (msfGFP) was inserted downstream of *acrB* on the chromosome to produce an AcrB-msfGFP fusion protein. This strain was created using the msfGFP from the pET GFP LIC cloning vector (u-msfGFP) which was a gift from Scott Gradia (Addgene plasmid # 29772; http://n2t.net/addgene:29772; RRID:Addgene_29772). Strain construction was based on the method used by Bergmiller et al. (2017) in *E. coli*^27^ where the codon optimised polylinker was used. Using restriction and ligation, the *aph* gene was inserted into pET LIC vector (u-msfGFP), so that strains containing the plasmid could be selected for. Using this plasmid as template, *gfp* and *aph* were inserted into the chromosome downstream of *acrB* in SL1344 to produce a protein fusion strain.

### Flow cytometry assay

The flow cytometric EtBr accumulation assay has been previously described^17^. Here this method was used to measure accumulation in samples from the same culture at different timepoints during batch culture. Briefly, cultures were grown at 37°C overnight in 5 mL of LB and sub-cultured at 4 % into fresh LB. A sample was taken at 0 hours and then every hour for 6 hours during growth. At each hour, sample volume was adjusted such that approximately 10^7^ cells were harvested and re-suspended in 1 x Hepes Buffered Saline (5X HBS; Alfa Aesar). Cells were washed and resuspended in 1 mL HBS. 100 μL of cell suspension was then further diluted into 500 μL HBS and SYTO™ 84 (Thermo Fisher Scientific) and ethidium bromide added to give final concentrations of 10 μM and 100 μM respectively. Samples were incubated for 10 minutes before measuring accumulation by flow cytometry. Flow cytometry settings and emission filters were used from Whittle et al^17^. Briefly, The SYTO 84 fluorescence emission was collected in the YL1-H channel (585/16 nm) using a 561 nm yellow laser and used to differentiate cells from acellular material.

EtBr fluorescence was collected using the BL3-H channel (695/40 nm) using a 488 nm blue laser. SYTO 84 accumulation measurements (**Figure 4**), was not a repeated experiment but data was re-analysed from EtBr accumulation assays and therefore fluorescence emission was collected in the YL1-H channel (585/16 nm) using a 561 nm yellow laser. Nile Red accumulation was measured as previously described^17^. In these experiments SYTO 9 (10 μM; Thermo Fisher Scientific) was used to differentiate cells from acellular particles using the BL2-H channel. Nile red has an excitation of 549 nm and emission of 628 nm in the presence of phospholipids, and in a neutral lipid environment (tryglycerides), the fluorescence shifts to ex/em of 510/580 nm^31^. Nile red fluorescence was excited using the yellow laser and detected using the YL1-H channel for orange fluorescence^17^.

### Flow cytometry assay in the presence of EDTA

Growing culture samples were taken at 1, 3 and 5 hours as above. Samples were made with varying concentrations of EDTA (0 μM, 1 μM, 10 μM, 100 μM, 200 μM and 500 μM) in 500 μL HBS. These concentrations of EDTA increased the final volume of the sample because the stock concentration was limited by solubility. Dyes were then added but volume added was adjusted to maintain the final concentration stated above. Once the dyes were added, 100 μL of cell suspension was added and cells were incubated for 10 minutes at room temperature. Samples were then analysed by flow cytometry.

### Whole population transcription analysis

Overnight cultures containing pMW82 transcriptional reporter plasmids were diluted 1:10000 in MOPs minimal media, supplemented with 50 μg/ml ampicillin. OD_600_ and GFP fluorescence were measured every 30 minutes for 12 hours using a Fluostar Omega (BMG labtech) incubated at 37 °C. OD_600_ and GFP fluorescence were measured, and a minimal media only control subtracted from the data. SL1344 autofluorescence was removed by subtracting SL1344 fluorescence from that of pMW82 strains. GFP fluorescence divided by OD_600_ was used as a measurement to disregard cell density across growth.

### Efflux assay

Efflux assays were carried out as previously described as previously^14^. This assay measures direct efflux activity of a population of cells by pre-loading cells with a fluorescent efflux substrate in the presence of the proton motive force inhibitor, CCCP, and re-energising cells with glucose to measure the decrease in fluorescence as substrates leave the cells. Briefly, overnight cultures of SL1344 and SL1344 Δ*acrB* were sub-cultured into fresh LB and then grown for 5 hours at 37°C. At the 1, 3 and 5-hour time points, 10mL of culture was taken and the OD_600_ measured. The harvested cell pellet was then resuspended in phosphate buffer containing MgCl_2_ buffer and each strain adjusted to the same OD_600_.

### RNAseq

The transcriptome of SL1344 and SL1344 Δ*acrB* were analysed at different time points during growth (1, 3 and 5 hours). There were 4 replicates of each strain. MOPS minimal media was inoculated at 4% with overnight cultures. Cultures were incubated at 37°C, shaking for 5 hours. At 1 hour, 5 mL of culture was centrifuged at 3500 x g for 5 minutes at room temperature to harvest the cells. The supernatant was removed and the pellet was snap frozen. At the 3 and 5-hour time points, only 1 mL of culture was harvested and snap frozen. GENEWIZ Inc. carried out the RNA extraction, quality control, library preparation, sequencing and bioinformatic analysis. Briefly, total RNA was extracted from *S*. Typhimurium cell pellets using RNeasy Plus Universal kit (Qiagen), and RNA quality control was carried out using Qubit 2.0 Fluorometer to measure total RNA concentration and Agilent TapeStation to produce an RNA integrity number (RIN) and a DV200 score. To remove rRNA, the ribozero Removal Kit was used (Illumina). The NEBNext Ultra II RNA Library Prep Kit (Illumina) was used for library preparation, following the manufacturer’s protocol. For library preparation, cDNA was synthesised, end repaired and adenylated at the 3’ ends. Universal adapters were ligated to cDNA and library enrichment was carried out using limited cycle PCR. Sequencing was carried out using Illumina HiSeq 4000. Bioinformatic data analysis was carried out by GENEWIZ Inc. Trimmed reads were mapped to the SL1344 reference genome FQ312003 using the Bowtie2 aligner. Unique gene hit counts were calculated by using feature Counts from the Subread package. All statistical analysis was performed using R. With the package, DESeq2, a comparison of gene expression between the groups of samples was performed. The Wald test was used to generate p-values and Log2 fold changes. Data is accessible on ArrayExpress with the accession code E-MTAB-9679.

## Supporting information

Supplementary information including methods, discussion and figures

## Authors Contributions Statement

JMAB, TWO and EEW designed these assays. GFP transcriptional reporter strains were constructed by ET. EEW and HM performed experiments to obtain samples for RNAseq. EEW performed all other experiments. EEW analysed all data. RNAseq data was analysed by EEW and TWO. This manuscript was written by EEW, JMAB, TWO and MAW.

