## Supplementary information including methods, discussion and figures for "Efflux only impacts drug accumulation in actively growing cells"

**Supplementary methods**

**Chromosomal insertion of *gfp* downstream of *acrB* to produce SL1344 AcrB-GFP**

To measure the protein level of AcrB in *S.* Typhimurium, a gene encoding a monomeric super‑folder GFP (msfGFP) was inserted downstream of *acrB* on the chromosome to produce a AcrB-msfGFP fusion protein. This strain was created using the msfGFP from the pET GFP LIC cloning vector (u-msfGFP) was a gift from Scott Gradia (Addgene plasmid # 29772 ; http://n2t.net/addgene:29772 ; RRID:Addgene_29772). Strain construction was based on the method used by Bergmiller et al. (2017) in *E. coli*^1^ where the codon optimised polylinker was used*.* Using restriction and ligation, the *aph* gene was inserted into pET LIC vector (u‑msfGFP), so that strains containing the plasmid could be selected for. Using this plasmid as template, *gfp* and *aph* were inserted into the chromosome downstream of *acrB* in SL1344 to produce a protein fusion strain.

The restriction sites downstream of the *gfp* gene that enable cutting by AscI and KpnI‑HF (NEB) restriction enzymes, were used to insert *aph* into the msfGFP encoding plasmid. The *aph* gene was amplified from the pKD4 plasmid. The forward oligonucleotide contained the AscI restriction site at the 5’ end, where the KpnI restriction site was present at the 5’ end of the reverse oligonucleotide. Restriction digest was carried out on the msfGFP plasmid and the *aph* DNA product using the enzymes stated above to produce overhangs of DNA for re‑ligation. The ligation reactions were then transformed in to NEB 5‑ α *E. coli* (C29871) using the NEB transformation protocol.

The msfGFP + *aph* plasmid was purified using the Qiagen plasmid prep protocol. Oligonucleotides were designed to amplify GFP and *aph* from the plasmid with homology to *acrB*. The forward oligonucleotide shown in **Table S1**, was designed to have homology to the 3’ end of *acrB,* a codon optimised polylinker (GgtAgcGgtAacAaaGgtCagGgc)^1^ and homology to the 5’ of *gfp.*The reverse oligonucleotide had homology to the end of *aph* and a non‑coding downstream region of *acrB*. Insertion of *gfp* and *aph* downstream of *acrB* was done using homologous recombination^2^,. PCR and sequencing confirmed correct fusion of *gfp* to *acrB* and then *aph* was removed using pCP20^2,3^.

| Table S1. List of oligonucleotides | | |
| --- | --- | --- |
| Code | **Description** | **Sequence (5’-3’)** |
| 242 | (F) for inserting *aph* into msfGFP with upstream AscI site | TATATTGGCGCGCCGTGTAGGCTGGAGCTGCTTC |
| 243 | (R) for inserting *aph* into msfGFP with downstream KpnI site | AGGATATTCATATGGACCATGGCTAATTCCCATGGTACCCCGATA |
| 246 | *aph* insertion into msfGFP (F) | GTCCAAGCTGAGCAAAGACC |
| 247 | *aph* insertion into msfGFP (R) | TGAATGAACTGCAGGACGAG |
| 261 | (F) to insert msfGFP-*aph* downstream of *acrB* | GAGCATAGTCATTCGACAGAACATCGCGGTAGCGGTAACAAAGGTCAGGGCGTGAGCAAGGGCGAGGAGCTGTT |
| 249 | (R) for the insertion of msfGFP-*aph* downstream of *acrB* | GGACCATGGCTAATTCCCATTTTGCTCACTGTTGATAAGGCCGCGCAAGCGGCCTTTTTTACGCAAAAATCT |

**Supplementary Discussion**

**Evidence for envelope remodeling from RNAseq**

Starting at the inner membrane, expression of *cfa* had significantly increased expression at 3 or 5 hours of growth compared to 1 hour in SL1344 (3.17-fold change and 7.27-fold change respectively). The *cfa* gene product converts linear unsaturated fatty acids (attached to phospholipids) to cyclopropane fatty acids via the addition of a methyl group donated by *S*-adenosylmethionine (SAM). It should also be noted that genes encoding the SAM synthesis pathway are highly upregulated at 3 and 5 hours of growth. It has previously been shown that cyclopropane fatty acids accumulate in the inner membrane in stationary phase, with the onset of starvation, and have been hypothesised to lead to a decrease in membrane fluidity^4–6^. It has also been shown that the *cfa* gene is in part regulated by RpoS^7,8^ Mutants lacking *cfa* display increased susceptibility to acid, heat and pressure in stationary phase, further highlighting the importance of cyclopropane fatty acids in tolerating these stresses via membrane mediated mechanisms^9,10^. It is therefore likely that increased *cfa* expression in stationary phase results in decreased inner membrane fluidity and enhanced barrier function.

***Peptidoglycan***

A key component of the Gram-negative envelope is the peptidoglycan (PG) cell wall. It has previously been shown that in stationary phase PG makes up a larger proportion of dry cell weight ^11^ and is more highly crosslinked^12^. Transpeptidase enzymes that introduce crosslinks between PG peptide chains were differentially expressed in our RNAseq dataset. In *E. coli*, the majority of peptidoglycan crosslinks are 4-3 or DD crosslinks between d-Ala and meso-diaminopimelic acid (mDAP). Of the 4 DD-transpeptidases, *mrcA* (PBP1A), *mrdA* (PBP2) and *ftsI* (PBP3) had decreased expression at 5 hours, and *mrcB* (PBP1B) remained unchanged. A smaller percentage of crosslinks occur between mDAP and mDAP; the number of these 3-3 or LD crosslinks increase dramatically in stationary phase^12–14^. Of the two LD-transpeptidases (LDTs) that form 3-3 crosslinks, transcription of *ldtD* was unchanged at 5 hours of growth compared to 1 hour in our dataset, but *ldtE* (*ynhG*) expression increased. In *E. coli*, *ldtE* has been described as the ‘housekeeping’ stationary phase LDT^13^; its expression is activated by RpoS and ppGpp^15,16^. It is proposed that 3-3 crosslinks reinforce the envelope in non-replicating conditions and in response to stress^17,18^. In clinical isolates, bacteria with increased LD crosslinks can be highly resistant to β-lactams because they do not require PBPs for transpeptidation^19^.

In *E. coli* three further LDTs (LdtA, LdtB and LdtC) have been shown to crosslink the PG to Braun lipoprotein Lpp, which links the PG to the OM and thus regulates the width of the periplasm^20,21^. The proportion of Lpp crosslinked in this manner increases in stationary phase^12^. In our dataset, *ldtA* was upregulated at 5 h while *ldtB* was downregulated at 3 and 5 hours. In *E. coli*, *ldtA* expression is RpoS-dependent^15^. A triple *E. coli* *ldtA ldtB ldtC* knockout leaks periplasmic proteins and is sensitive to EDTA, as is an *lpp* mutant, although a double *ldtD ldtE* knockout is not^17^. It is therefore suggested that the PG-OM linkage via Lpp is critical for OM stability and barrier function^19^. It is also worth noting that *lppB* expression increased at 5 h in our dataset; *S.* Typhimurium has two genes encoding Lpp, *lppA* and *lppB*.

Two genes encoding glycosyltransferases that polymerise PG from Lipid II subunits were oppositely regulated in our dataset; *mrcA* (PBP1A) was downregulated and *mtgA* upregulated at 3 and 5h. We believe that this observation is novel.

Expression of PG hydrolases was also seen to change in our RNASeq data; this class of enzymes is essential for PG turnover and remodelling, cell growth, and other functions^22^. Three DD-endopeptidases (MepS, MepM and PbpG) which break crosslinks between d-Ala and mDAP, had decreased expression at 3h and 5h. These endopeptidases are required for PG remodeling to permit cell elongation during exponential growth^23^.

Four genes encoding lytic transglycosylases (*mltA, mltC, mltD*, and *mltF*) that cleave bonds between the sugar residues are downregulated at 5 h. This class of enzymes are important for multiple functions including PG remodeling during growth^24,25^, so it is logical that they are downregulated in stationary phase. It has been suggested that their contribution to cleavage and cell division is reduced in comparison to amidases in *E. coli*^26^.

The DD-carboxyopetidases *dacA* (PBP5) and *dacC* (PBP6) which cleave the terminal d-Ala from PG peptide chains were observed to be down- and up-regulated at 5 h in our dataset. Expression of *dacC* is induced in stationary phase and is ppGpp activated^16^. DacC is postulated to play a role in stationary phase PG stabilization whereas DacA is thought to aid regulation of crosslinking and maintenance of cell shape in exponential phase^22^. Similar to *dacA* and *dacC*, two genes encoding amidases *amiC* and *amiD* which cleave the pentapeptide chain from the sugar backbone of PG were oppositely regulated in our dataset. AmiC in *E. coli* plays the largest role in PG degradation and remodelling during cell division and septation^26^. The role of AmiD is poorly understood although our data suggest a novel role in stationary phase.

Taken together, stationary phase gene expression leads to more LD-crosslinks and more crosslinks to the OM via LPP, both contributing to decreasing envelope permeability.

***Outer membrane***

The phospholipid composition of the inner leaflet of the OM is known to change in stationary phase, with an increased cardiolipin concentration required for viability in *E. coli*^27^*.* Cardiolipin synthase B (*clsB*) was upregulated at 5 h in our dataset.

Genes involved in lipid A biosynthesis had decreased expression at 5 hours, suggesting that stationary phase lipid A production is downregulated, likely due to decreased growth. An increase in LPS in stationary phase has previously been linked to increased cell death^28^.

Most genes involved in lipid A modification that were significantly altered in this dataset had decreased expression at 5 hours compared to 1 hour. These include genes responsible for resistance to cationic antimicrobial peptides including polymyxin^29^. The addition of 4-amino-4-deoxy-L-arabinose (l-Ara4N; *arn* genes) and ethanolamine (*eptA*) to the phosphate groups of lipid A reduces its net negative charge^29^. Similar negative charge-reducing modifications occur in the oligosaccharide core of LPS^30^ and have an impact on CAMP resistance^31^.

Palmitate incorporation into lipid A by PagP also confers resistance to CAMPs (although not polymyxin B) by increasing OM hydrophobicity. The PmrAB two-component regulatory system is the key regulator of LPS modification and polymyxin resistance^29^ and is also downregulated at 5h in our dataset. It therefore appears that these LPS modifications are more prevalent in exponential growth and not required in stationary phase. Finally, a number of genes encoding O-antigen synthesis enzymes are downregulated at 5h; average O-antigen chain length is known to increase in stationary phase^32^ and O-antigen structure has been shown to influence serum resistance^32^ and CAMP susceptibility^33^.

Only one lipid A modification gene was upregulated at 5 hours: *lpxO* encodes a dioxygenase that hydroxylates a myristoyl chain in lipid A^34^. Deletion of *lpxO* in *S*. Typhimurium decreases survival in macrophages^35^ and in *Klebsiella pneumonia*, an *lpxO* mutant is more sensitive to CAMPs^36^.

Enterobacterial common antigen (ECA) when linked to LPS or in its cyclic form has been linked to envelope integrity and bile resistance^37–39^. Six genes involved in the biosynthesis of ECA had decreased expression at 5h. It has previously been shown ECA is not converted to cyclic ECA in mutants lacking WecA, WecF or WecG^40^, and as expression of the *wecG* decreased, it may be deduced that there is less cyclic ECA and less ECA in general in stationary phase.

***Regulatory networks***

Supplementary Table S1 also outlines the known regulation of genes differentially regulated in our RNASeq dataset, either in *Salmonella* or *E. coli*. Our EtBr accumulation data in Fig 5 suggested that RpoS only plays a minor role in envelope remodeling giving rise to decreased EtBr influx. In contrast, RpoS is seen to play a major role in envelope remodeling leading to increased SDS resistance in *E. coli*^41^; however, this points to differences in regulation of envelope remodeling leading to different resistance phenotypes. The increase in CAMP resistance of stationary-phase *Salmonella* is RpoS-independent and instead partially dependent on PhoPQ^42^. Indeed, PhoPQ has been shown to be a key regulator of envelope barrier function^43^; influx of three dyes into stationary phase *S.* Typhimurium was greater in the absence than the presence of PhoP. The PmrAB regulon was seen to decrease in expression at 5 h. PmrA coordinates LPS modifications to reduce sensitivity to CAMPs^44^, but our data suggests that these modifications are far more important in exponential phase, aligning with our ‘division of labour’ model in stationary phase whereby each layer of the envelope plays a larger role in barrier function.

**Figure S1 Transcription of efflux pumps across growth using GFP-transcription reporters**

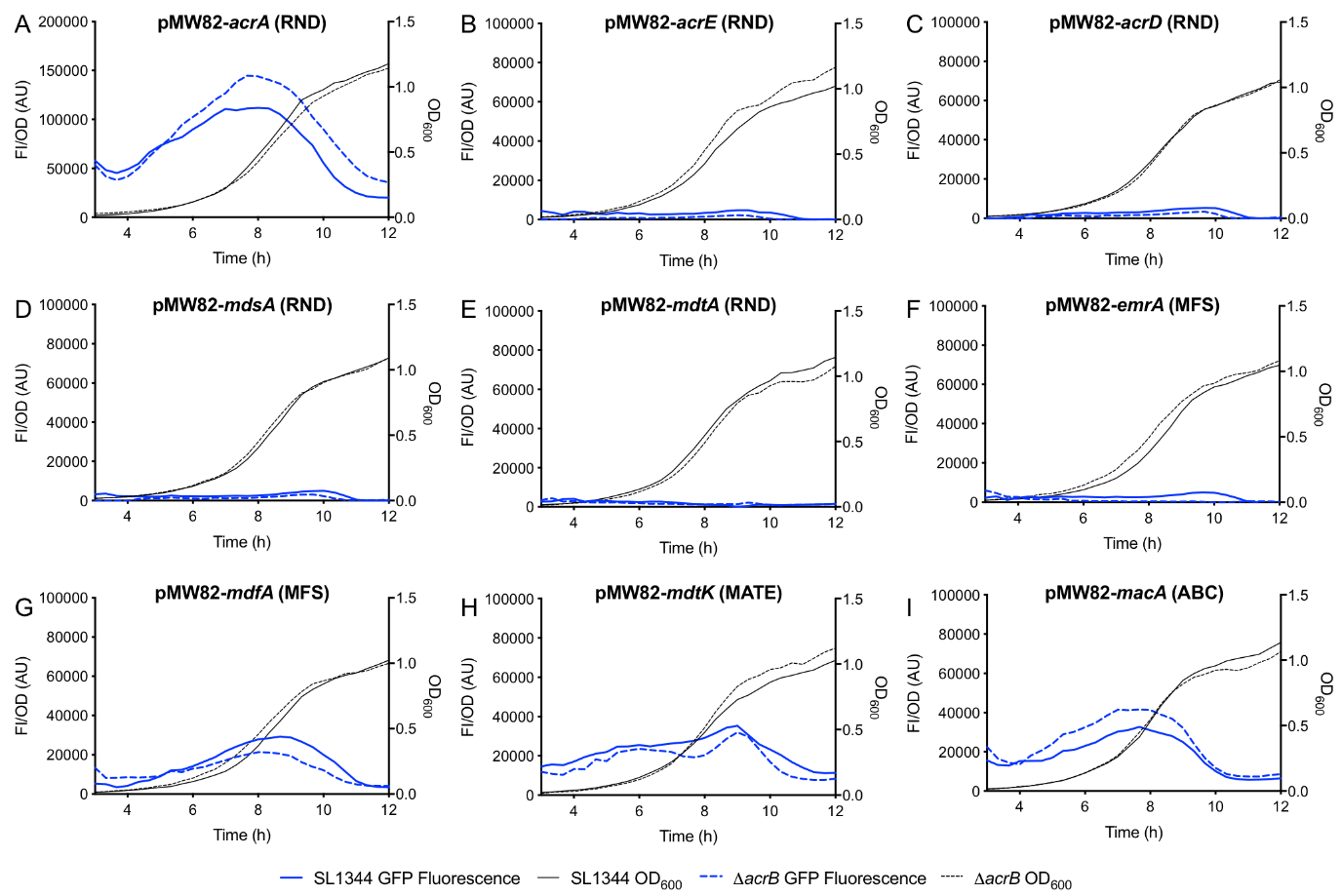

In the following graphs **A‑I**, the GFP/OD_600_ (measured using a plate reader) and therefore transcription of each pump is represented as a solid blue line (SL1344) or a dashed blue line (∆acrB). The OD_600_ was also plotted and shown as a solid black line (SL1344) or a dashed black line (∆acrB). Each graph represents the fluorescence of a different transcriptional reporter: (**A**) acrA, (**B**) acrE, (**C**) acrD, (**E**) mdsA, (**D**) mdtA, (**F**) emrA, (**G**) mdfA, (**H**) mdtK and (**I**) macA.

**Figure S2 EtBr accumulation in SL1344 + 100 µM CCCP**
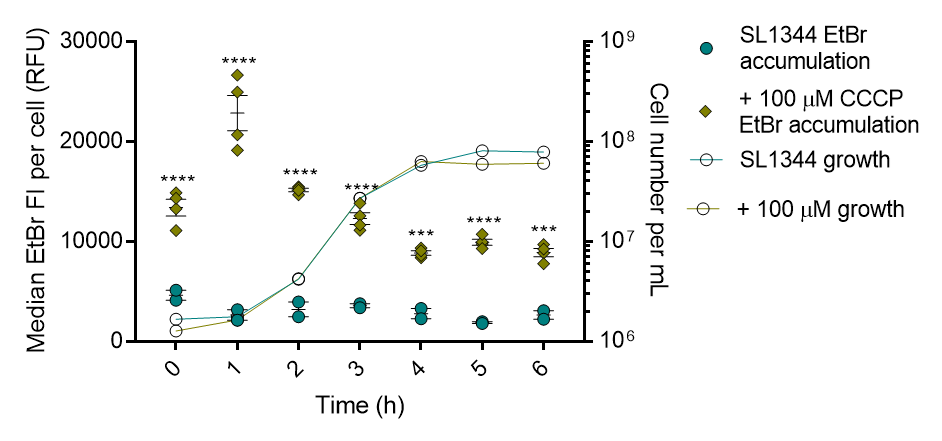

SL1344 vs SL1344 + 100 µM CCCP**.** Median EtBr fluorescence per cell in 10,000 SYTO‑84^+^ flow cytometry events was measured every hour between 0 and 6 hours. Individual green circles (WT) and individual green diamonds (+ CCCP) represent the median value of EtBr fluorescence within a biological replicate. 2 biological replicates for the control strains are shown, and 4 replicates for those with CCCP, with a short mean bar and SEM error bars. EtBr accumulation is plotted on the left Y‑axis. Calculated cell number values were plotted on the right Y‑axis with corresponding symbols equating to strain and a dashed line to show growth of the culture. Cell numbers were based on the mean of the same biological replicates and the same gated population that EtBr fluorescence was measured from. Two‑way ANOVA and Sidak’s multiple comparisons test were used for statistical analysis; At 0, 1, 2, 3 and 5 hours, EtBr accumulation was significantly increased in the presence of CCCP with p values of <0.0001 (****). At 4 hours and 6 hours, EtBr accumulation was significantly increased with p values of 0.0003 (***) and 0.0002 (***) respectively.

**Figure S3 EtBr accumulation in E. coli and K. pneumoniae efflux mutants**

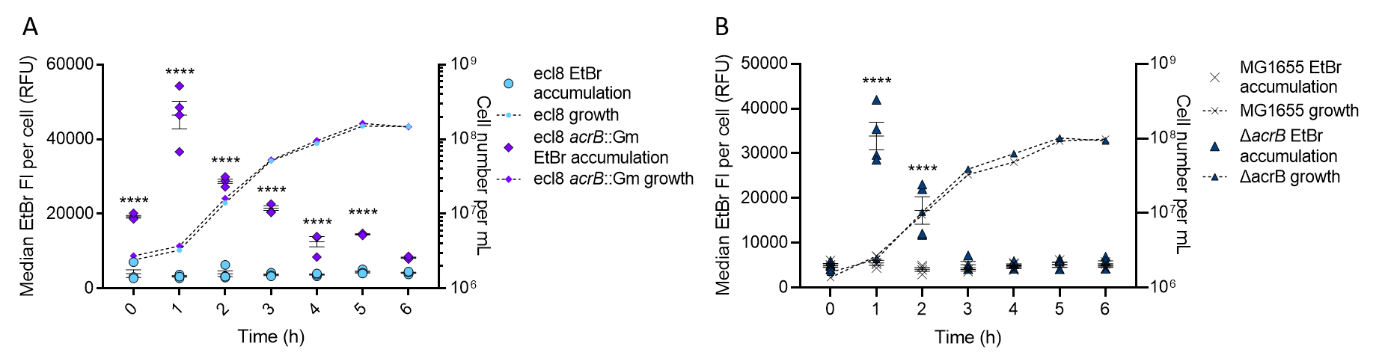

**(A)** shows K. pneumoniae ecl8 and ecl8 acrB::Gm. Median EtBr fluorescence per cell in 10,000 SYTO‑84^+^ flow cytometry events were measured every hour between 0 and 6 hours. Individual blue circles (WT) and individual purple diamonds (acrB::Gm) represent the median value of EtBr fluorescence within a biological replicate. **(B)** shows E. coli MG1655 and ΔacrB**.** Median EtBr fluorescence per cell in 10,000 SYTO‑84^+^ flow cytometry events was measured every hour between 0 and 6 hours. Individual black Xs (WT) and individual blue triangles (ΔacrB) represent the median value of EtBr fluorescence within a biological replicate. 4 biological replicates for each strain are shown, with a short mean bar and SEM error bars. EtBr accumulation is plotted on the left Y‑axis. Calculated cell number values were plotted on the right Y‑axis with corresponding symbols equating to strain and a dashed line to show growth of the culture. Cell numbers were based on the mean of the same biological replicates and the same gated population that EtBr fluorescence was measured from. Two‑way ANOVA and Sidak’s multiple comparison test were carried out for statistical analysis. In K. pneumoniae, EtBr accumulation is significantly increased in acrB::Gm at 0, 1, 2, 3, 4 and 5 hours with p values of <0.0001 (****). In E. coli, EtBr accumulation is significantly increased in ΔacrB at 1 and 2 hours with p values of <0.0001 (****).

**Figure S4 EtBr accumulation in *P. aeruginosa***

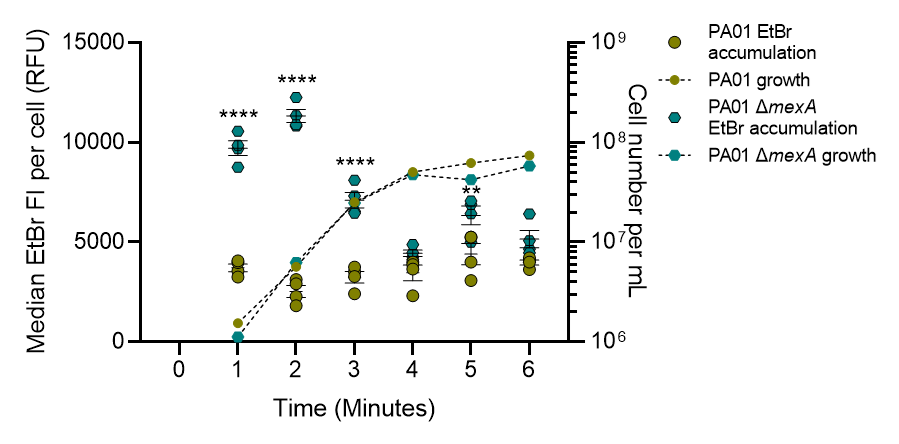

Median EtBr fluorescence per cell in 10,000 SYTO‑84^+^ flow cytometry events was measured every hour between 0 and 6 hours. Individual green circles (WT) and individual blue hexagons (ΔmexA) represent the median value of EtBr fluorescence within a biological replicate.4 biological replicates for each strain are shown, with a short mean bar and SEM error bars. EtBr accumulation is plotted on the left Y‑axis. Calculated cell number values were plotted on the right Y‑axis with corresponding symbols equating to strain and a dashed line to show growth of the culture. Cell numbers were based on the mean of the same biological replicates and the same gated population that EtBr fluorescence was measured from. At 1, 2 and 3 hours, EtBr accumulation is significantly increased in ΔmexA with p values of <0.0001 (****) and at 5 hours, a p value of 0.0024 (**).

**Figure S5 EtBr accumulation in SL1344 and SL1344 Δ*acrB* grown in MOPs minimal media**

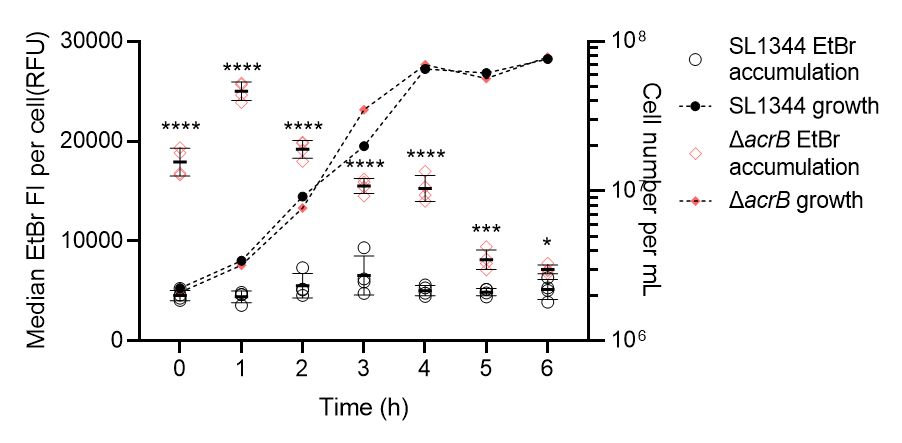

Median EtBr fluorescence per cell in 10,000 SYTO‑84^+^ flow cytometry events was measured every hour between 0 and 6 hours. Individual black circles represent the X‑median value of EtBr fluorescence in SL1344 (WT) (from 10,000 SYTO‑84^+^ cell events) within a biological replicate, therefore the average accumulation of ethidium per cell. Individual pink diamonds represent the X‑median value of EtBr fluorescence in SL1344 ΔacrB (from 10,000 SYTO‑84^+^ cell events) within a biological replicate. 4 biological replicates for each strain are shown, with a short mean bar and SEM error bars. EtBr accumulation is plotted on the left Y‑axis. Calculated cell number values were plotted on the right Y‑axis with corresponding symbols equating to strain and a dashed line to show growth of the culture. Cell numbers were based on the mean of the same biological replicates and the same gated population that EtBr fluorescence was measured from. Two‑way ANOVA and Sidak’s multiple comparison test were carried out for statistical analysis. P values at 0, 1, 2, 3 and 4 hours was <0.0001 (****), at 5 hours was 0.0003 (***) and at 6 hours was 0.0494 (*).

**Figure S6 Nile Red accumulation in SL1344 and SL1344 Δ*tolC***

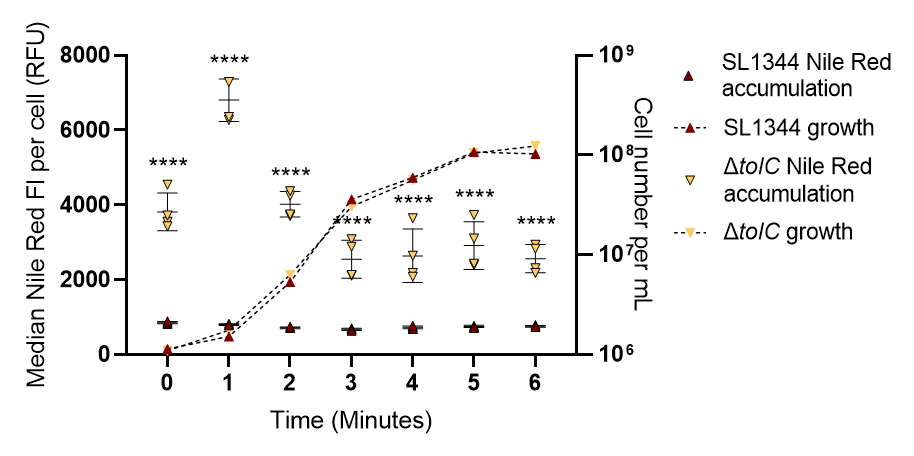

Median Nile Red fluorescence per cell in 10,000 SYTO‑9^+^ flow cytometry events was measured every hour between 1 and 6 hours. 0 hour time points were not included as SYTO‑9^+^ populations could not be gated. Single red triangles represent the X‑median value of nile red fluorescence in SL1344 (WT) (from 10,000 SYTO‑9^+^ cell events) within a biological replicate. Individual orange triangles represent the X‑median nile red fluorescence in ΔtolC (from 10,000 SYTO‑9^+^ cell events) within a biological replicate. 4 biological replicates for each strain are shown, with a short mean bar and SEM error bars. Nile red accumulation is plotted on the left Y‑axis. Calculated cell number values were plotted on the right Y‑axis with corresponding symbols equating to strain and a dashed line to show growth of the culture. Cell numbers were based on the mean of the same biological replicates and the same gated population that nile red fluorescence was measured from. Two‑way ANOVA and Sidak’s multiple comparison test were carried out for statistical analysis and ‘****’ related to P <0.0001.

**Figure S7 EtBr accumulation in porin deleted strains of SL1344 and SL1344 Δ*acrB***

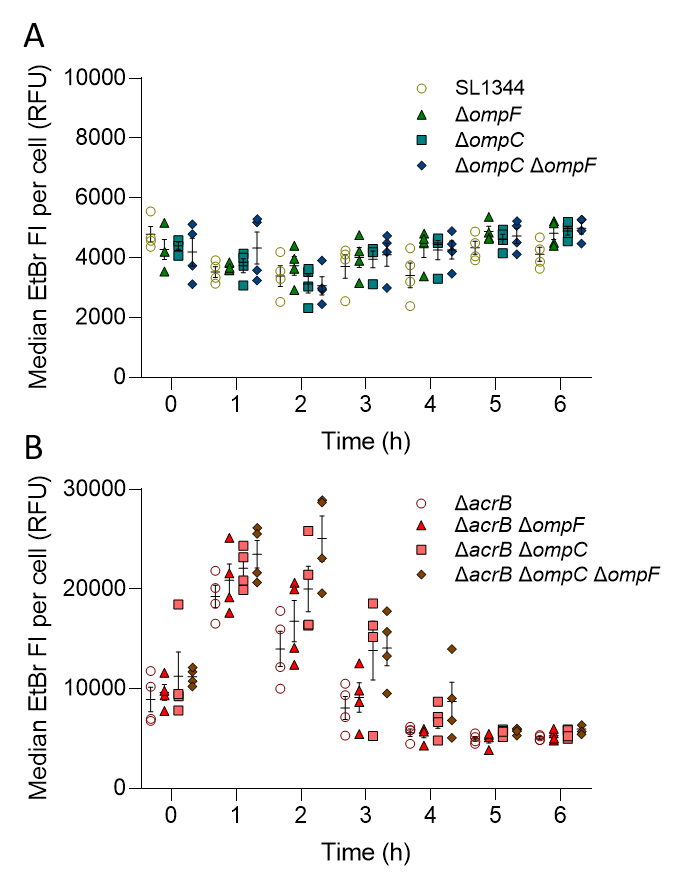

**(A+B)** 4 biological replicates for each strain are shown, with a short mean bar and SEM error bars. EtBr accumulation is plotted on the left Y‑axis **(A)** shows SL1344 WT (individual green dots) vs ΔompF (green triangles), ΔompC (green square) and ΔompC ΔompF (blue diamonds)**.** Median EtBr fluorescence per cell in 10,000 SYTO‑84^+^ flow cytometry events was measured every hour between 0 and 6 hours. Individual symbols represent the median value of EtBr fluorescence within a biological replicate. **(B)** shows SL1344 ΔacrB (red dots) vs ΔacrB ΔompF (red triangles), ΔacrB ΔompC (pink square) and ΔacrB ΔompC ΔompF (red diamonds)**.** Median EtBr fluorescence per cell in 10,000 SYTO‑84^+^ flow cytometry events was measured every hour between 0 and 6 hours. Individual symbols represent the median value of EtBr fluorescence within a biological replicate. A two‑way ANOVA and Dunnett’s multiple comparison test were used for statistical analysis.

**Table S1.** Genes implicated in envelope remodeling that were identified by RNASeq as being differentially expressed between 1 and 3 or 1 and 5 hours growth.

| **Pathways** | **Function** | | **gene** | **1 vs 3 hr fold change** | **1 vs 5 hr fold change** | **Regulation** |
| --- | --- | --- | --- | --- | --- | --- |
| CFA biosynthesis | Conversion of unsaturated fatty acids to cyclopropane fatty acids | | *cfa* | 3.17 | 7.27 | Ec – RpoS and ppGpp activated, sRNA regulated |
| L-methionine and SAM biosynthesis | Synthesis of the methyl donor SAM | | *metJ* | 4.29 | 2.08 |  |
|  |  |  | *metQ* | 4.33 | 2.30 | Ec – MetJ repressed |
|  |  |  | *metC* | 10.71 | 4.15 | Ec – MetJ repressed |
|  |  |  | *metK* | 11.02 | 6.20 | Ec – MetJ repressed |
|  |  |  | *metI* | 12.00 | 4.36 | Ec – MetJ repressed |
|  |  |  | *metN* | 17.43 | 3.79 | Ec – MetJ repressed |
|  |  |  | *metL* | 27.63 | 3.20 | Ec – MetJ repressed, PhoP activated |
|  |  |  | *metA* | 42.82 | 13.61 | Ec – MetJ repressed |
|  |  |  | *metB* | 48.08 | 2.21 | Ec – MetJ repressed, PhoP activated |
|  |  |  | *metR* | 51.96 | 5.54 | Ec – MetJ, MetR repressed |
|  |  |  | *metF* | 155.57 | 37.62 | Ec – MetJ repressed |
|  |  |  | *metE* | 236.20 | 64.66 | Ec – MetJ repressed, MetR activated |
| Peptidoglycan biosynthesis and remodelling | glycosyltransferases | | *mrcA* | 0.40 | 0.29 |  |
|  |  |  | *mtgA* | 2.41 | 2.88 |  |
|  | D,D‑carboxypeptidases | PG remodelling | *mrcA* | 0.40 | 0.29 |  |
|  |  |  | *dacA* | 0.46 | 0.19 |  |
|  |  |  | *dacC* | - | 2.34 | Ec – ppGpp activated |
|  | D,D‑transpeptidases | Forms 4-3 PGcrosslinks | *mrcA* | 0.40 | 0.29 |  |
|  |  |  | *mrdA* | 0.47 | 0.29 |  |
|  |  |  | *ftsI* | 0.29 | 0.45 |  |
|  | L,D‑transpeptidases | Crosslinks PG to Lpp | *ldtA* | - | 2.50 | Ec – RpoS activated |
|  |  |  | *ldtB* | 0.12 | 0.10 |  |
|  |  | Forms 3-3 PG crosslinks | *ldtE* | 2.44 | 2.96 | Ec – RpoS and ppGpp activated |
|  | Amidases | PG remodelling | *amiC* | 0.49 | 0.46 |  |
|  |  |  | *amiD (ybjR)* | - | 2.41 |  |
|  | Lytic transglycosylases | PG remodelling | *mltA* | - | 0.26 |  |
|  |  |  | *mltC* | - | 0.35 |  |
|  |  |  | *mltD* | 0.30 | 0.11 |  |
|  |  |  | *mltF* | 0.38 | 0.39 |  |
|  | D,D‑endopeptidases | PG turnover and remodelling | *mepS* | 0.31 | 0.25 |  |
|  |  |  | *mepM* | 0.31 | 0.26 |  |
|  |  |  | *pbpG* | 0.36 | 0.36 |  |
| Braun lipoprotein | Links OM to PG | | *lppB* |  | 2.10 |  |
| Cardiolipin synthesis | Synthesis of cardiolipin from phosphatidylglycerol | | *clpB* |  | 2.61 | Ec - ppGpp activated |
| Lipid A biosynthesis | Synthesis of lipid A component of LPS | | *lpxH* | 0.28 | 0.17 |  |
|  |  |  | *lpxB* | 0.41 | 0.21 |  |
|  |  |  | *lpxK* | 0.38 | 0.21 |  |
|  |  |  | *lpxA* | - | 0.25 |  |
|  |  |  | *lpxL* | - | 0.30 |  |
|  |  |  | *lpxD* | - | 0.41 |  |
| Lipid A modifications | Two‑component regulatory system, regulator of polymyxin resistance | | *pmrA* | - | 0.18 |  |
|  |  |  | *pmrB* | 0.49 | 0.11 |  |
|  | Addition of 4-amino-4-deoxy-L-arabinose (L‑Ara4N) to lipid A | | *arnA (pmrI)* | - | 0.07 | ST- PmrAB activated |
|  |  |  | *arnB (pmrH)* | - | 0.06 | ST- PmrAB activated |
|  |  |  | *arnC (pmrF)* | - | 0.09 | ST- PmrAB activated |
|  |  |  | *udg (pmrE)* | - | 0.10 | ST- PmrAB activated |
|  |  |  | *arnT (pmrK)* | 0.46 | - | ST- PmrAB activated |
|  | Myristoyl chain hydroxylation | | *lpxO* | - | 2.54 | ST- PhoPQ activated |
|  | Addition of ethanolamine to lipid A | | *eptA (pmrC)* | - | 0.06 | ST- PmrAB activated |
|  | Addition of palmitate to lipid A and phospholipids | | *pagP* | 0.28 | 0.11 | Ec – PhoP activated |
|  | Palmitoleote Incorporation | | *lpxP* | 0.13 | 0.05 |  |
| LPS core modifications | Adds phosphoethanolamine to heptose (I) phosphate in LPS core | | *cptA* | - | 0.19 | ST- PmrAB activated |
|  | Dephosphorylates heptose(II) phosphate in LPS core | | *pmrG (ais)* | - | 0.08 | ST - PmrAB activated |
| O antigen synthesis and modification | O-antigen acyltransferase | | *oafA* | - | 0.08 |  |
|  | O-antigen translocase | | *wzx (rfbX)* | - | 0.49 |  |
|  | O-antigen polymerase | | *wzy (rfc)* | - | 0.39 |  |
|  | O antigen biosynthesis rhamnosyltransferase | | *wbaN (rfbN)* | - | 0.35 |  |
|  | O-antigen chain length determinant protein Wzz_ST_ | | *wzzB* | 0.39 | 0.18 | ST- PmrAB activated |
|  | O-antigen ligase | | *waaL (rfaL)* | 0.28 | 0.16 |  |
| ECA subunit biosynthesis | UDP-N-acetyl-D-mannosamine dehydrogenase | | *wecC* | - | 0.40 |  |
|  | dTDP-glucose 4,6-dehydratase | | *rffG* |  | 0.43 |  |
|  | dTDP-fucosamine acetyltransferase | | *wecD* | - | 0.39 |  |
|  | Lipid III flippase | | *wzxE* | 0.45 | 0.34 |  |
|  | TDP-N-acetylfucosamine:lipid II N-acetylfucosaminyltransferase | | *wecF* | 0.39 | 0.21 |  |
|  | UDP-N-acetyl-D-mannosaminuronic acid transferase | | *wecG* | - | 0.47 |  |

Fold changes are shown as 3h/1h or 5h/1h so that a positive value indicates higher expression at 3h or 5h. A dash signifies no significant difference in expression observed at that timepoint. Regulation data for *E. coli* (Ec) and *Salmonella* Typhimurium (ST) are taken from Ecocyc and the literature cited in the manuscript.

6.	Huisman, G. W., Siegele, D. a, Zambrano, M. M. & Kolter, R. Morphological and physiological changes during stationary phase. *Escherichia coli Salmonella Cell. Mol. Biol.* (1996).
